# Soil microbiome functions in plant-aphid dynamics are resilient to community restructuring during compost application

**DOI:** 10.64898/2026.10.04.756575

**Authors:** Mary M. Dixon, Ethan J. McAnally, Brian Chang, Elias H. Bloom, Clare L. Casteel

## Abstract

**Background:** Soil microbiomes play critical roles in plant health and resilience, yet it is still poorly understood how disruptions in local microbial communities, such as during soil management, impact these ecological functions. In this study, we investigated how compost addition, a widely used soil management practice, and microbiome origin influence microbiome-mediated plant resistance to insects. We combined surveys of working organic farms across New York with controlled laboratory experiments, amplicon sequencing, plant-insect bioassays, and mass spectrometry.

**Results:** Compost use was associated with shifts in the relative abundance of 20 fungal and 2 bacterial genera across ten working farms. However, no consistent changes in bacterial or fungal diversity or community structure in fields with compost were observed across the farms. Under controlled conditions using a common compost source, compost increased rhizosphere microbial diversity and significantly altered rhizosphere community structure, including the relative abundance of 6 fungal and 41 bacterial genera. Despite these substantial community-level changes, compost did not affect populations of green peach aphids (*Myzus persicae*) on kale (*Brassica oleracea*). Instead, aphid populations and plant salicylic acid concentrations were more strongly associated with microbiome origin, demonstrating that compost can substantially restructure soil and rhizosphere microbial communities without corresponding changes in microbiome-mediated plant resistance to insects.

**Conclusions:** Our findings demonstrate that microbial diversity and compositional responses to management are not necessarily reliable indicators of all microbiome functions. More broadly, this work highlights the importance of local soil microbiome variation in shaping plant-insect interactions and emphasizes the need to link community-level responses to direct functional measurements when evaluating microbiome responses to agricultural management.

**Highlights:** Soil microbiome origin impacts plant-aphid dynamics and salicylic acid levels. While compost application increases microbial diversity and homogenized community structure, it did not alter farm-specific effects of the microbiome on plant-aphid interactions.

## Background

It is well established that soil microbiome composition plays a critical role in regulating plant health and performance [1–3]. For example, plant growth promoting rhizobacteria (PGPR) can support a range of beneficial functions for the host plant [4] including enhanced nutrient acquisition [5,6], improved tolerance to drought stress [7], and suppression of soil-borne pathogens [8]. Beyond their belowground functions, PGPR can also stimulate induced systemic resistance (ISR) [9], a plant-wide defense state that helps protect aboveground tissue from some pathogens and insect herbivores [10]. Accumulating evidence suggests that components of the soil microbiome beyond PGPR can also influence plant resistance to insect herbivores [11,12], although the specific microbial players remain largely unknown. While these beneficial microbes offer promising avenues for improving crop productivity and soil health, their assembly and functions can be highly context dependent [13], limiting broad translation across agroecosystems.

Agricultural practices are used to shape the soil microbiome in predictable ways, with important consequences for plant defense and plant-insect interactions. Long-term organic farms are associated with decreased pest populations, increased soil microbiome diversity [14,15], and greater abundance of plant-beneficial rhizosphere microbes [16]. In a few cases, shifts in soil microbiome composition on organic farms have been directly linked to enhanced plant defense signaling [14] and nematode suppressive soil microbiomes [17]. For instance, lower leafhopper incidence was observed in organically managed tomato (*Solanum lycopersicum*) fields compared to paired conventional tomato fields [14]. Using microbiome transfer experiments, laboratory bioassays, and tomato signaling mutants, lower leafhopper preferences was linked to changes in the microbiome and induction of salicylic acid (SA), a phytohormone regulator of plant defense [14]. Despite organic systems have been associated with reduced pest pressures and more diverse soil microbiomes, significant variation in outcomes have been observed. For example in the study above, there was no decrease in leafhopper preference for one of the organic farms compared to the paired conventional sites but there was still an increase soil microbiome alpha diversity [14], suggesting other microbiome or farm characteristics are mediating to pest suppression outcomes.

Compost is widely used in organic agriculture to provide nutrition to crops and promote soil microbiome health [18,19] through the introduction of microbial taxa and nutritional resources, increased abundance of plant-beneficial microbes [20], and microbiome network complexity [21]. However, the outcomes of compost on plant-insect interactions are variable. In some cases compost addition is associated with increased pest susceptibility, such as through increased aphid (*Myzus persicae*) populations in bean (*Phaseolus vulgaris*) [22]. In contrast, compost diminished woolly apple aphid populations (*Eriosoma lanigerum*) in apple (*Malus domestica*) [23] and brown planthoppers (*Nilaparvata lugens*) in rice (*Oryza sativa*) [24]. Differences in starting soil microbiome community structure have been shown to influence the impact of agronomic interventions on other ecological outcomes. For example, tillage and herbicide application decreased microbial decomposition rates on farms in Germany, while the same treatments increased microbial decomposition on farms in France [25]. Despite results indicating that specific organic management promotes natural pest management through soil microbiome change, it remains unclear how different soil microbiomes respond to agricultural practices and disruptions.

In this study we hypothesized that compost application would restructure soil microbiome composition, but its effects on microbiome-mediated functions in plant defense would depend on the starting soil microbiome structure. To test this hypothesis, we leveraged natural variation in compost use across working organic vegetable farms in New York and controlled laboratory experiments with field-collected soils. In the field survey, amplicon sequencing was used to characterize fungal and bacterial communities associated with compost application. To disentangle the effects of compost and soil microbiome origin from variations in environmental factors and management, we then transferred microbiomes from different farms to a common soil background in laboratory experiments and amended each with a single compost source. In these experiments kale (*Brassica oleracea*), a crop with increasing total production in organic agriculture [26,27], and *Myzus persicae*, an agriculturally important pest and virus vector [28], were used as a model plant-insect system to assess how compost application and microbiome origin influence microbiome-mediated functions in plant-insect interactions.

## Methods

### Field trial and Illumina sequencing data

In 2021, we surveyed 279 organic vegetable farmers on practice use and microbiome beliefs [19]. Farmers were also asked to submit soil samples for sequencing from two different field plots where specific agronomic practices had been used consistently for at least 1 year [29]. Out of the 85 farmers that submitted soil samples, 10 farmers submitted two samples from a composted field and an un-composted field that were managed otherwise similarly (Fig. 2A). The published soil metabarcoding data from the V4-V5 region of the bacterial 16S rRNA gene and the fungal ITS2 region were obtained for the paired samples with and without compost for each of the 10 farms (NCBI BioProject Accession: PRJNA1334013). These data had expected read number and quality (Table S1) [29]. Read preprocessing, data clustering, and post-clustering were conducted in AMPtk [25][30], taxonomy was assigned using the SILVA version 138.1 [31] and UNITE version 29.11.2022 [32] databases for bacteria and fungi, respectfully using DADA2 in R [33]. Relative abundance of specific bacterial and fungal taxa were analyzed between composted and un-composted sites using rarified OTU count tables and the ANCOM-BC2 package in R [34]. A prevalence threshold of 0.10 was used with an = 0.01 and a Holm p adjustment method.

Alpha diversity measures (Shannon’s diversity, OTU richness) were calculated using the phyloseq package [35] and with rarified count tables for bacterial and fungal taxa between the paired composted sites and un-composted sites. Repeated measures ANOVA were used to evaluate differences (compost vs no compost) with farm as a random effect: Y ∼ Compost + (1|Farm); Y = diversity measure (Shannon, OTU richness) (n = 10 replicates of compost and un-composted soil, N = 20). Model assumptions were evaluated using diagnostic plots [36]. Beta diversity analysis was performed by calculating Bray-Curtis distance on Hellinger-transformed relative abundance data and ordinated using principal coordinate analysis (PCoA) with the vegan package [37]. Differences in community composition between composted and un-composted samples were tested using PERMANOVA with the adonis2 function (permutations constrained by farm to reflect the paired design), while multivariate dispersion was assessed with the betadisper function using the vegan package in R [37]. A differential abundance analysis was conducted with the log-fold change in abundance of taxa in the composted compared to the un-composted treatments for the lab experiments using the ANCOMBC2 function [34] as outlined above. For this analysis, a log fold change cutoff of 1.5 was used, as well as a prevalence threshold of 0.10 with α = 0.01 and a Holm p adjustment.

### Soil collection for laboratory experiments

To isolate impacts of compost application on soil microbe communities from other variables in the field we collected soil from four organic farms across New York and conducted lab experiments. In April 2023 and in 2024, bulk soil was collected in a transect across the field to a 15 cm depth and combined for each site for that year. Farming practices used at each site were collected in an abbreviated survey, as in [29]. After collection, soil was immediately stored at 4 °C until use. In 2023 a portion of the soil was used for DNA extractions and bioassays, while soil was only for DNA extractions in 2024.

### Bulk soil DNA extractions, Nanopore sequencing, and bioinformatic analysis

DNA was extracted form a portion of the field collected soil using the Qiagen DNeasy PowerSoil Pro Kit (Qiagen, Hilden, Germany), quantified, and cleaned as previously described [29]. Takara PrimeStar polymerase (Takara Bio, Kusatsu, Shiga, Japan) was used to amplify the 18S rRNA region (NS1-F and Fun-rOP-R [38]) and the 16S-23S rRNA region (16S-rRNA-27-F and 23S-rRNA-2241-R [29,39].]). For 18S amplification, reactions were run with 30 ng template with an initial cycle of 3 min at 94 °C, 25 cycles (10 s at 98 °C, 15 s at 60 °C, 4.5 min at 68 °C), and final elongation for 7 min 68 °C. For 16S-23S amplification, 15 ng template was used, and reactions occurred with an initial cycle for 3 min at 94 °C, 25 cycles (10 s 98 °C, 15 s 60 °C, 4 min 68 °C), and a final cycle of 7 min at 68 °C. Following PCR, samples were cleaned with Monarch Spin PCR & DNA Cleanup Kit (New England Biolabs, Ipswich, MA) as described above and concentrations quantified with a Qubit fluorometer (Thermo Fisher Scientific, Waltham, MA) using the High Sensitivity dsDNA assay kit (Thermo Fisher Scientific, Waltham, MA). Samples were pooled into a library at equal concentration to a total quantity of 500 fmol in 50 μL. Library preparation and sequencing was done with the Oxford Nanopore SQK-LSK114 kit (Oxford Nanopore Technologies, Oxford, UK) following manufacturer instructions. Oxford Nanopore sequencing was used for several reasons including its ability to produce longer reads with greater resolution and accuracy for microbial communities [14].

BAM files of reads were filtered to duplex only reads and converted to a single FASTQ file using SAMtools [40]. Duplex basecalling was run using Oxford Nanopore Dorado software set to the high Accuracy mode. Reads were filtered by length and quality using cutadapt [41]. Cutadapt was also used for demultiplexing, retaining and labelling only reads with both a forward and reverse barcode region. Read quality and length distribution were assessed after filtering to confirm retention of high-quality reads. Read abundance from each sample was quantified, and samples were evenly rarified by randomly selecting 3000 reads from each sample. Clustering of reads to OTUs was performed at 97 % identity using UPARSE within the AMPtk pathway [34]. [42]. Singleton OTUs were discarded. Taxonomy was assigned using the DADA2 assigntaxonomy command in R [33]. Fungal reads were aligned to the UNITE+INSD eukaryote dataset [32], and bacterial reads were aligned to the RDP dataset at the species level [32,43]. Fungal reads not aligning to the fungal kingdom were removed from the dataset. Read length and quality scores were consistent across sequenced libraries (Table S2). Relative abundance of phyla was calculated across the two replicates collected from the four field sites (4 farms x 2 years = 8 samples) and beta diversity analysis was performed by calculating Bray-Curtis distance on Hellinger-transformed relative abundance data and ordinated using principal coordinate analysis (PCoA) using the vegan package [37].

### Microbiome transfers in the laboratory, compost application, and plant growth conditions

The remaining soil after sequencing was used in microbiome transfers experiments. Briefly, 16 g of the field collected soil was suspended in 240 mL of quarter strength Hoagland’s solution, shaken at 275 rpm for 1 h at room temperature and then allowed to settle for one hour before transferring the supernatant to 250 mL centrifuge bottles. The supernatant was then centrifuged at 500 rpm for 5 minutes at 4 °C and stored for up to 2 weeks before use in experiments. Two kale seeds (*Brassica oleracea,* ‘Red Russian’) were placed in 5 cm × 5 cm-cell planting trays filled with Lambert LM111 mix (Lambert Peat Moss Inc., Rivière-Ouelle, Québec, Canada) that had been autoclaved 3 times for 30 minutes at 121 °C and 262 kPa. Each week, 15 mL of each soil microbiome extract treatment was added to a separate tray of plants. Half of all treatments were amended with 30 % non-sterile commercial compost (Premium Cayuga Compost, Cayuga Compost, Trumansburg, NY) one week after planting. Each treatment was bottom-watered two times a week and fertilized with a quarter-strength Miracle-Gro All Purpose Plant Food (N-P_2_O_5_-K_2_O: 24-8-16). Plants were thinned to one plant per cell after two weeks.

### Insect colonies, lab bioassays, leaf collection, and rhizosphere soil collection

Green peach aphids (*Myzus persicae*) were reared at 25 °C in 16:8 L:D cycle as described previously [44]. After 4 weeks, 2 adult aphids were placed in a clip cage on the fourth true leaf on half of the experimental plants. Twenty four hours later both adults and all but one nymph were removed to ensure experimental insects were the same age. Empty cages were added to the remaining plants as a control. After nine days the foundress had developed into a reproducing adult and the resulting population in each cage was counted. Samples were removed if the initial foundress escaped or the leaf broke off during experimentation. Each plant was also gently removed from the pots, bulk soil gently removed from roots, and rhizosphere soil collected as described before [14]. At the end of the experiment, the caged leaves without aphids were cut from each plant and leaf tissue immediately frozen in liquid nitrogen and stored at -80 °C until salicylic acid extractions.

### Salicylic acid measures

Our previous work demonstrated that tomato plants grown on in soil microbiomes from long term organic farms have increased salicylic acid (SA) levels compared to plants grown in conventional microbiomes [14]. To determine whether variation in microbiome-mediated SA accumulation is associated with changes in plant–aphid interactions in the present study, we next quantified SA levels in the kale leaf tissue using liquid chromatography–mass spectrometry (LC–MS). Briefly, leaf tissue was lyophilized, weighed, and approximately 10 mg of dried tissue was homogenized and then extracted in a buffer spike with an isotopically labeled internal standard for salicylic acid (D4-SA), as in [14,29]. Dried extracts were resuspended in 200 μL HPLC-grade methanol, and 10 μL was injected onto an Dionex UHPLC (Thermo Fisher Scientific, Waltham, MA, USA) equipped with a C18 reversed-phase column (Phenomenex Gemini) and coupled to an Orbitrap-Q Exactive mass spectrometer (Thermo Scientific, Waltham, MA, USA) operated in negative-ionization mode. Compounds were separated using a gradient of 0.1 % (v/v) formic acid in water and 0.1 % (v/v) formic acid in acetonitrile at a flow rate of 600 μL min^−1^. Peak areas were quantified using Xcalibur (Thermo Fisher Scientific, Waltham, MA, USA), and endogenous SA abundance was normalized to the D4-SA internal standard and sample dry weight.

### Statistical analysis of aphid bioassay and salicylic acid data

All statistical analyses were conducted in R (version 4.5.2) [45] and RStudio (version 2026.5.0.218) [46]. Impacts of compost treatment and soil microbiome origin on aphid fecundity and salicylic acid were assessed using an ANOVA (Y ∼ Compost * Farm). Assumptions were checked using diagnostic plots, and no transformations were necessary to satisfy assumptions [36]. Post-hoc pairwise comparisons were conducted using Fisher’s Least Significant Difference (LSD) test, as it is commonly used in aphid bioassay experiments [47–49], for differences between compost treatments and microbiome treatment using the agricolae package [50]. A Tukey HSD test was used to examine pairwise comparisons for the SA data. At least seven replicates for each treatment were used for the aphid bioassay statistical analysis (4 farms x 2 compost treatments x 7 replicates = 46) and five replicates were used for salicylic acid analysis (4 farms x 5 replicates = 20). Replicates were omitted from aphid fecundity analysis when the initial aphid escaped or the petiole of the leaf with aphid broke during manipulation.

### Rhizosphere microbiome characterization in the laboratory experiments

A subset of rhizosphere samples was selected from the laboratory experiment to evaluate impacts of compost on microbiome communities. For these experiments we selected the Farm 1 (red) and Farm 2 (blue) based on differences geographic location and starting microbiome structure, with the goal of capturing the most diversity. Farms were assigned numbers and colors to protect their identity. DNA was extracted, nanopore libraries prepared, sequencing performed, and the same bioinformatic pipeline used as described above. Shannon’s diversity index and OTU richness were calculated using rarefied count data with the phyloseq package [35]. A two-way ANOVA (Y ∼ Compost * Farm) was used to test for impacts of compost treatment and microbiome origin on either OTU Richness or Shannon’s diversity using the car package [51], with Tukey Honest Standard Difference (HSD) for post-hoc testing using the agricolae package [50]. To examine changes in microbiome structure, we calculated Bray-Curtis distance matrices calculated from Hellinger-transformed data using the vegan package [37]. Calculations were based on the relative abundance of microbial taxa and differences were visualized using Principal Coordinate Analysis (PCoA) with the cmdscale function from the R base stats package [45]. Differences in microbial community composition among treatments were assessed using Permutational Multivariate Analysis of Variance (PERMANOVA) based on Bray-Curtis distance matrices (Bray-Curtis Distance ∼ Compost * Farm). The beta dispersion of the microbial community was tested for Compost and Farm with an ANOVA using the car package [51] (Y ∼ Compost * Farm).

## Results

### Compost-associated shifts in soil microbiome structure are farm dependent

In our previous study [29], ten farms submitted soil samples from fields with and without compost application that were otherwise managed similarly (Fig. 1A, B; N = 10 farms x 2 fields each = 20 soil samples). Across these samples, 9,343 bacterial and 6,740 fungal operational taxonomic units (OTUs) were identified, representing 565 bacterial and 593 fungal genera, respectively. Principal coordinates analysis (PCoA) based on Bray–Curtis dissimilarities revealed that no conserved shifts in bacterial and fungal community structure were associated with compost application across farms (Fig. 1C; p < 0.05). Some farms showed relatively small differences in microbial community structure between compost-amended and non-amended fields (e.g., Farms G, C, J, and H), whereas others showed large shifts in community structure (e.g., Farms E and F) (Fig. 1C). Although farms spanned multiple regions of New York and differed somewhat in other production practices (Fig. 1A, B), neither geographic location nor these management differences corresponded to clear clustering patterns in the ordination observationally (Fig. 1C).

**Fig. 1.**
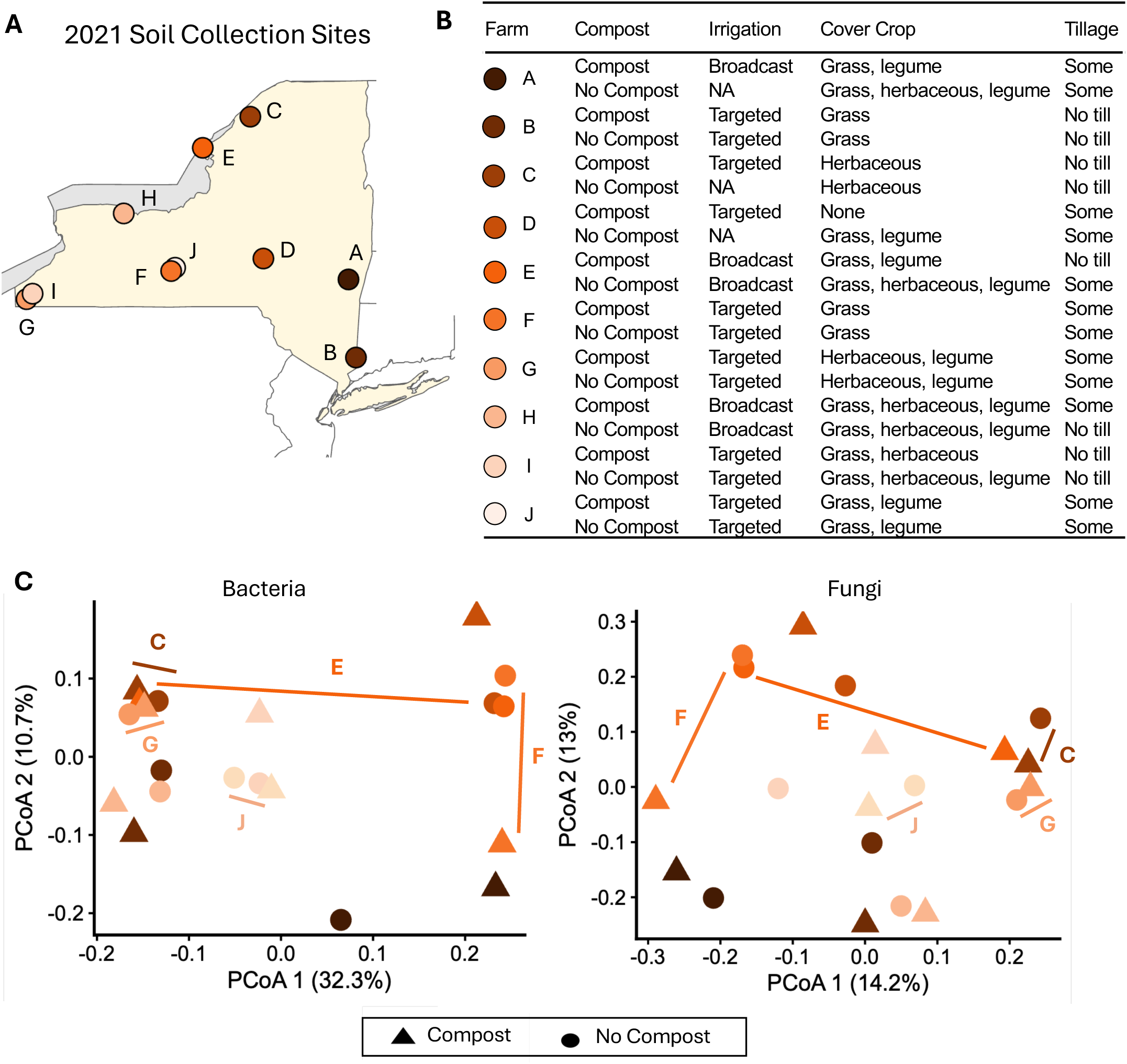
Compost-associated shifts in soil microbiome structure are farm dependent. (A) Location of the ten organic vegetable farms that submitted soil samples from fields with and without compost for amplicon sequencing in Bloom et al. (2026). (B) A description of management practices used on each field for the last three years that were reported by each farmer. (C) Beta diversity was ordinated with a PCoA with Bray Curtis for bacteria and fungi. Letters and lines represent the paired samples from the farms and are used to highlight differences.

### Compost application shifts specific soil taxa across working organic farms in New York

Compost application was not associated with significant changes in Shannon’s diversity (Fig. 2A; *p* > 0.05) or OTU richness (Fig. S1; *p* > 0.05) for bacterial or fungal communities. However, differential abundance analysis identified a subset of bacterial and fungal taxa that responded consistently to compost application across farms (Fig. 2B; Table S3; *p* < 0.05; log FC > 1.5). A greater number of fungal taxa (20) than bacterial taxa (2) changed in soils where compost was applied (Fig. 2B; Table S3). Among bacterial genera, *Escherichia-Shigella* (log FC = 1.95) and *Leptolyngbya* (log FC = 1.82), both increased in relative abundance with compost application. Twelve fungal genera increased and eight decreased in relative abundance with compost. The largest increases were observed for *Phialocephala* (log FC = 4.14), *Arachniotus* (log FC = 3.19), and *Hydeomyces* (log FC = 3.05), while the strongest decreases were observed for *Wickerhamomyces* (log FC = 1.66), *Spermospora* (log FC = 2.22), and *Steccherinum* (log FC = 2.10) (Fig. 2B, Table S3). Taken together these results demonstrates while overall soil microbiome diversity and structure are not impacted across diverse farms, compost application has conserved impacts on individual fungal genera, and to a lesser extent bacterial genera.

**Fig. 2.**
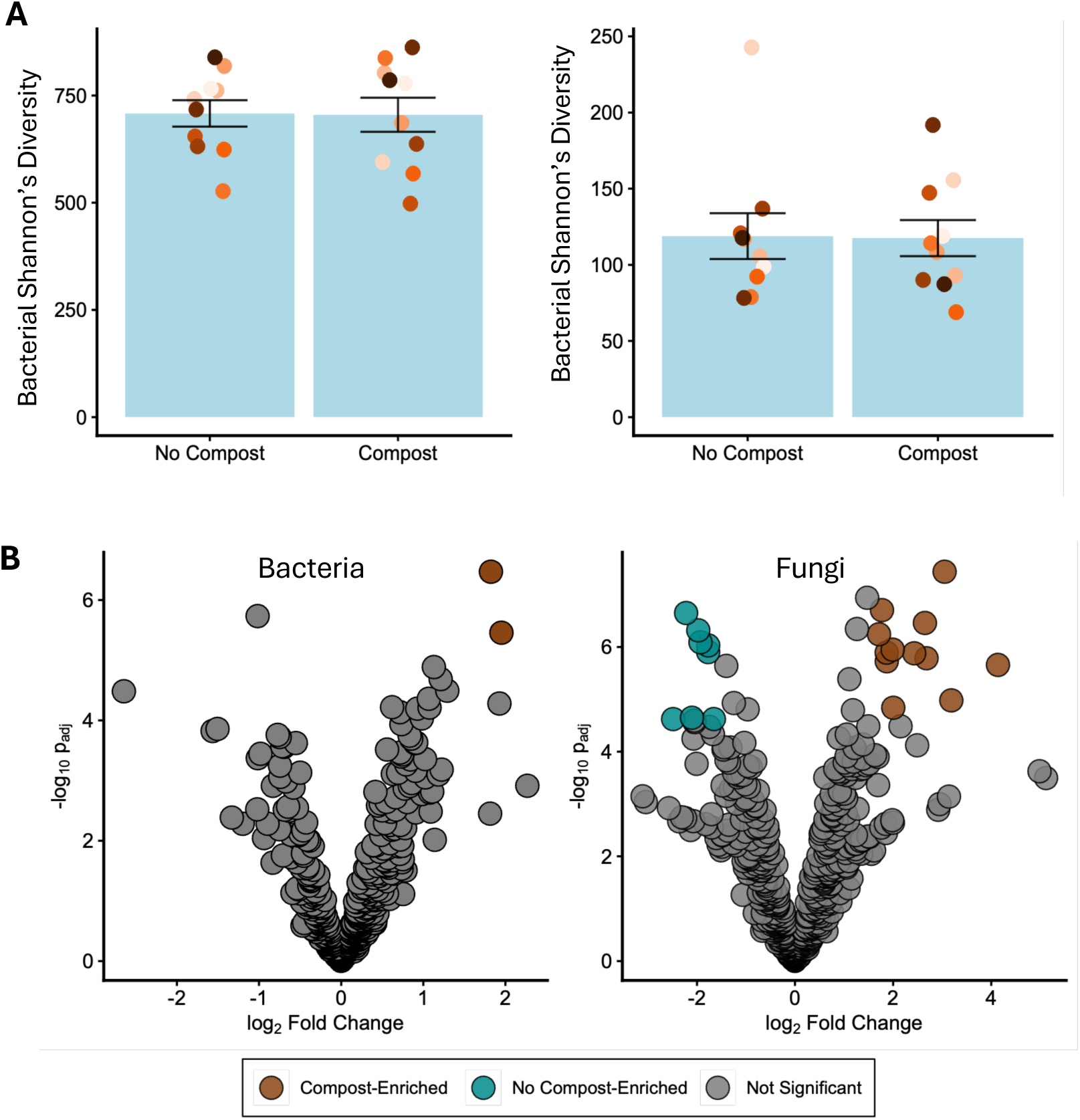
Compost application shifts specific taxa across working organic farms in New York. (A) Alpha diversity was captured by Shannon’s diversity for bacteria and fungi. Bars represent mean ± SE. (B) A volcano plot was used to illustrate the differentially abundant bacterial and fungal genera in field where compost was applied compared to fields without compost. Significant differences were determined using ANCOMBC with holm-adjusted-p < 0.05 and a log fold change > 1.5. Blue represents taxa that decreased and brown represents taxa that increased with compost application. Those taxa that did not significantly change are colored gray.

### Compost application shifts specific soil taxa across organic farms in New York

Compost application was not associated with significant changes in Shannon’s diversity (Fig. 2A; *p* > 0.05) or OTU richness (Fig. S1) in bacterial or fungal communities. However, differential abundance analysis identified a subset of bacterial and fungal taxa that responded consistently to compost application across farms (Fig. 2B; Table S1). Overall, substantially more fungal than bacterial taxa showed consistent changes in relative abundance in compost-amended soils (20 and 2 genera, respectively; Fig. 2B; Table S3; *p* < 0.05; |log FC| > 1.5). Among bacteria, *Escherichia-Shigella* (log FC = 1.95) and *Leptolyngbya* (log FC = 1.82) were consistently enriched across sites in response to compost application (Fig. 2b; Table S3). Twelve fungal genera increased in relative abundance in compost-amended fields, with the largest increases observed for *Phialocephala* (log FC = 4.14), *Arachniotus* (log FC = 3.19), and *Hydeomyces* (log FC = 3.05). Compost application was also associated with decreased relative abundance of eight fungal genera, with the largest decreases observed for *Wickerhamomyces* (log FC = −2.49), *Spermospora* (log FC = −2.22), and *Steccherinum* (log FC = −2.10) (Fig. 2B; Table S3). These results demonstrate that although compost application did not alter overall microbial diversity, it was associated with consistent shifts in the relative abundance of specific fungal genera and, to a lesser extent, bacterial genera across farms.

### Soil microbial communities differ across organic field sites used in lab studies

To examine the effects of compost and soil microbiome origin in more detail and separate their impacts from other environmental factors, we collected soil in 2023 and 2024 from four farms located in geographically distinct areas for additional experiments (Fig. 3A). Across farms and years, we identified 1,798 bacterial and 810 fungal OTUs. Ordination of bacterial and fungal communities showed that samples from the same farm generally clustered more closely than samples from different farms, regardless of collection year (Fig. 3B). Because each farm was represented by a single bulked sample per year, these patterns are descriptive and farm and year effects were not statistically tested. Taxonomic composition also varied among farms, with approximately 17 bacterial and 13 fungal phyla detected across the four sites (Fig. 3C). Across farms, the most abundant bacterial phyla were Actinomycetota, followed by Pseudomonadota and Bacillota, whereas the most abundant fungal phyla were Ascomycota, Mortierellomycota, and Basidiomycota (Fig. 3C).

**Fig. 3.**
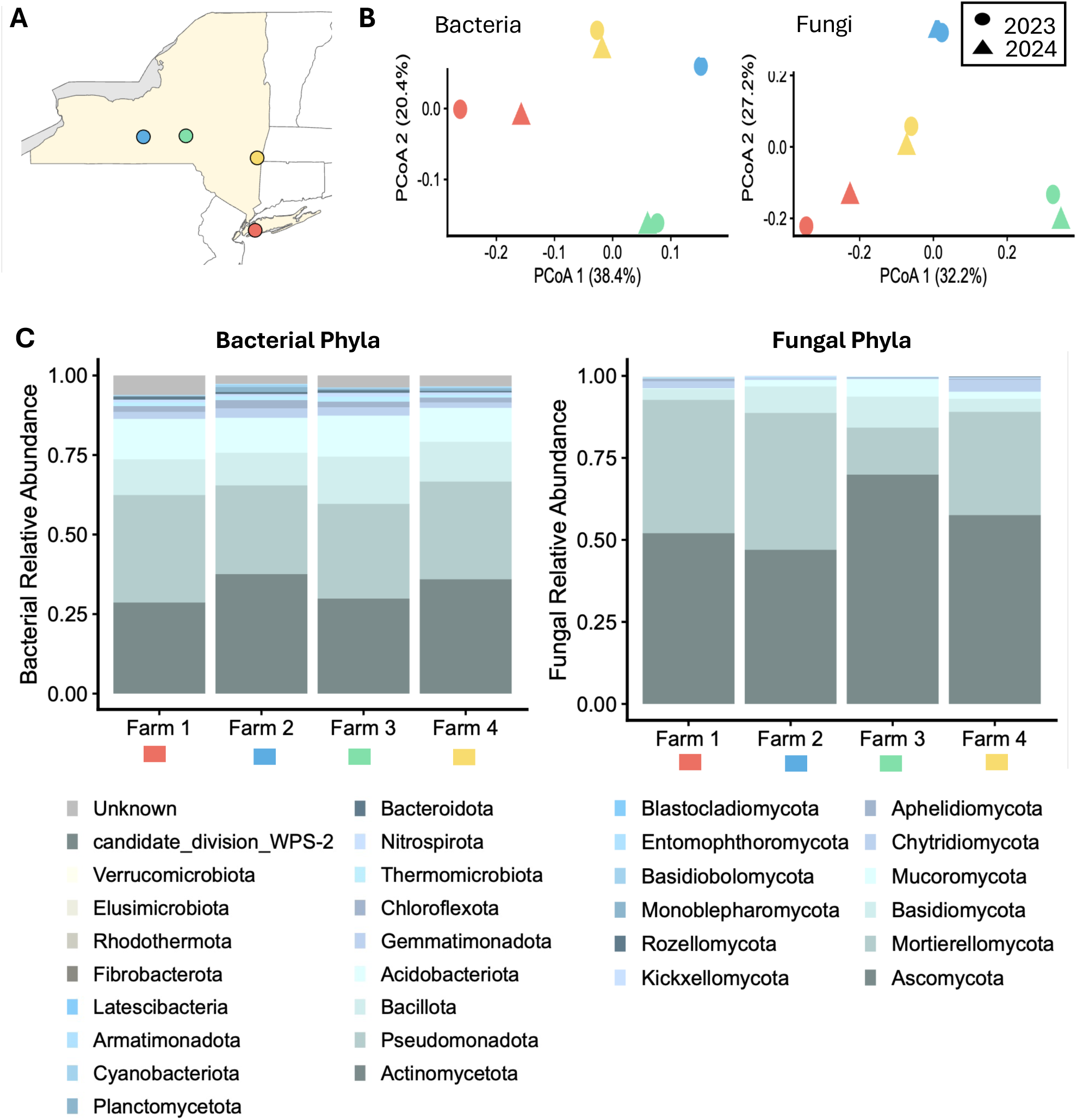
Soil microbial communities differ across organic field sites used in lab studies. (A) Soil samples were collected from four new samples across NY state in 2023 and 2024 and soil microbial communities were characterized using amplicon sequencing. (B) Beta diversity of bacterial and fungal communities were visualized in a PCoA ordination using Bray-Curtis distance. Color denotes the farm origin of the soil used: red (Farm 1 farm), blue (Farm 2), green (Farm 3), and yellow (Farm 4). Phyla was determined for each bacterial and fungal OTUs and relative abundance calculated for each farm across both years (C). OTUs that did not reach a phylum-level classification when summed by phyla for each cluster and included in the “Unknown” classification. N = 8.

### Microbiome origin, but not compost application, affects aphid populations and plant defense

To isolate differences in soil microbiome composition from differences in soil physical properties, microbial communities from each farm sample were transferred to a common soil substrate with and without compost and used to grow kale. In aphid bioassays, compost application had no significant effect on aphid populations on plants (Fig. 4A; compost main effect, *p* > 0.05). In contrast, soil microbiome origin significantly affected aphid populations (Fig. 4A; microbiome origin main effect, *p* = 0.02). The highest aphid populations were observed on kale plants grown in the soil microbiome from Farm 2 (Fig. 4A; blue farm), while the fewest aphids were observed on plants grown with the Farm 4 microbiome (Fig. 4A; yellow farm). Aphid populations did not differ between plants grown in microbiomes from Farm 2 (blue farm) and Farm 1 (red farm), or between plants grown with microbiomes from Farm 3 (green farm) and Farm 4 (yellow farm) (Fig. 4A). There was no significant interaction between microbiome origin and compost application (*p* = 0.22).

**Fig. 4.**
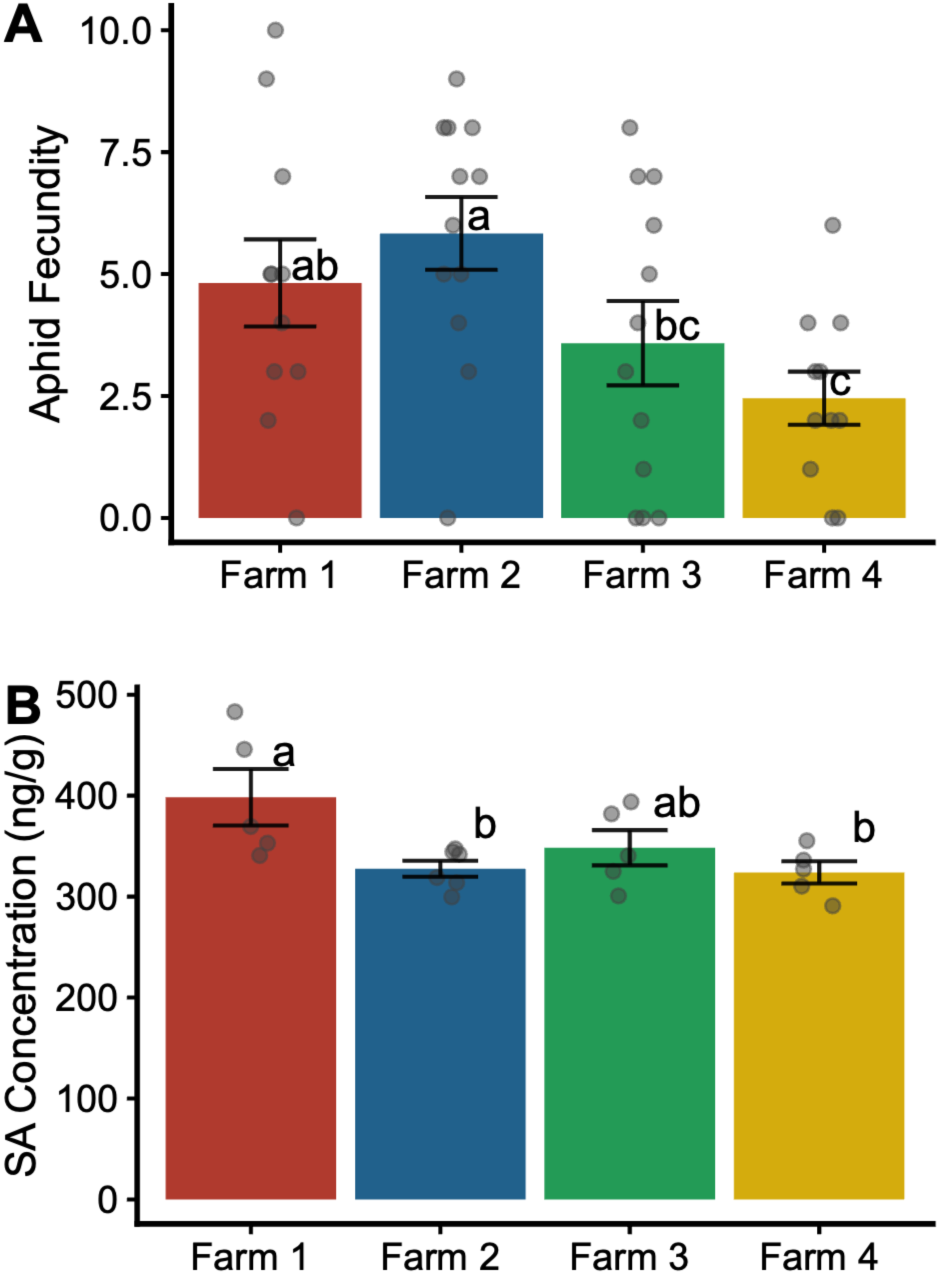
Microbiome origin, but not compost application, affects aphid populations and plant defense signaling. Soil microbiome transplants were performed in the lab using soil collected from different organic vegetable farms in New York. Kale (Brassica oleracea) was grown in the different transplanted soil microbiomes with and without compost addition for four weeks and then a single one-day-old Myzus persicae aphid added. (A) Nine days later aphid populations were measured. Aphid fecundity was calculated (N = 46) (B) and salicylic acid concentrations measured in a separate set of undamaged plants (N = 20). An ANOVA was run with Tukey HSD for pairwise comparisons. No significant main effects or interaction for compost were observed so results are presented at the farm level where there were differences. Different letters denote differences at α = 0.05 for the farm site used for soil collection. Bars represent mean ± SE. Color denotes the farm origin of the soil used: red (Farm 1), blue (Farm 2), green (Farm 3), and yellow (Farm 4).

To investigate whether variation in plant defense signaling was associated with microbiome-dependent variation in aphid populations, we quantified salicylic acid (SA) in plants grown with each soil microbiome in the absence of aphids. Consistent with the aphid population results, soil microbiome source significantly affected plant SA concentrations (Fig. 4B; *p* = 0.03). Kale plants grown in the Farm 1 microbiome (red farm) had the highest SA concentrations, while plants grown with microbiomes from Farm 4 (yellow farm) and Farm 2 (blue farm) had significantly lower SA concentrations (Fig. 4B). Although aphid populations did not differ significantly between plants grown with the Farm 1 and Farm 2 microbiomes (Fig. 4A), SA concentrations differed between these treatments (Fig. 4B), suggesting that additional plant or microbial factors contribute to microbiome-dependent effects on aphid populations.

### Compost application increases rhizosphere microbial diversity and alters community structure

We next characterized rhizosphere microbial communities from a subset of the bioassay plants to determine whether compost application altered microbial communities despite having no detectable effect on aphid populations. We selected microbiomes from Farms 1 and 2 because they differed in initial community structure (Fig. 3B) and induced different SA concentrations in kale (Fig. 4B), despite producing similar aphid populations (Fig. 4A). Compost application significantly increased bacterial and fungal Shannon’s diversity (Fig. 5A, B; bacterial, *p* = 3.63 × 10⁻¹⁰; fungal, *p* = 4.35 × 10⁻⁴) and OTU richness (Fig. S2A; bacterial, *p* = 4.31 × 10^-8^; fungal, *p* = 1.17 × 10^-7^). Bacterial diversity also differed between microbiome sources (Fig. 5A; microbiome origin main effect, *p* = 0.01), with higher diversity in rhizospheres derived from Farm 2 (blue) than Farm 1 (red). Fungal diversity did not differ between microbiome sources (Fig. 5B; microbiome origin main effect, *p* = 0.76). There was no significant compost x microbiome origin interaction for bacterial or fungal Shannon’s diversity (Fig. 5A, B; *p* > 0.05) and OTU richness was not significantly affected by microbiome origin or the compost x microbiome origin interaction (Fig. S2A; *p* > 0.05).

**Fig. 5.**
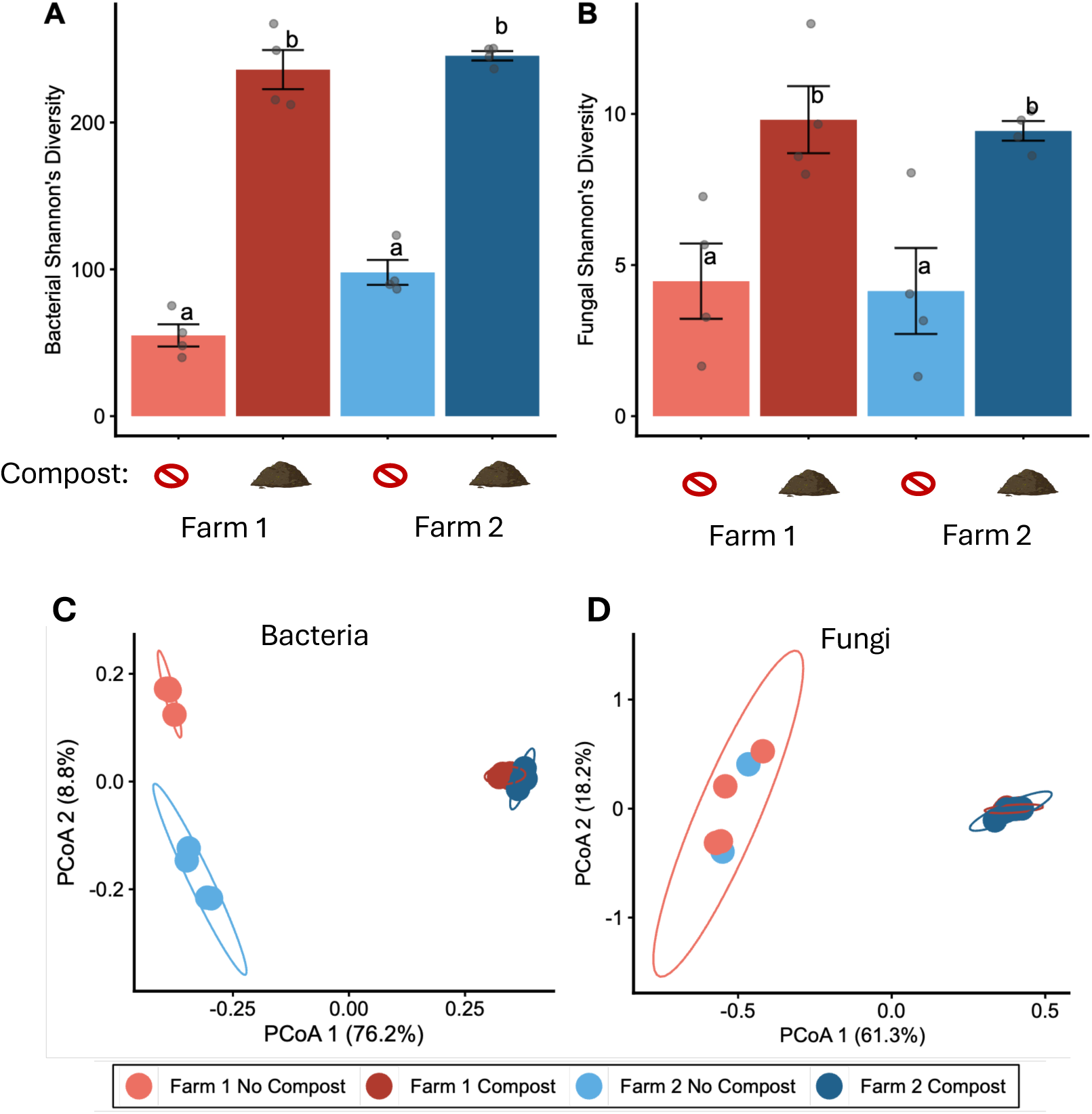
Compost application increases rhizosphere microbial diversity and alters community structure. Alpha diversity was captured by Shannon’s diversity for (A) bacteria and (B) fungi. Beta diversity was ordinated with a PCoA using Bray-Curtis distance for (C) bacteria and (D) fungi. An ANOVA was run with Tukey HSD for pairwise comparisons. Different letters denote differences at α = 0.05 for the farm site used for soil collection. Bars represent mean ± SE. Color denotes the farm origin of the soil used and the compost application: light red and dark red (Farm 1) & light blue and dark blue (Farm 2). N = 16.

Principal coordinates analysis (PCoA) revealed that rhizosphere bacterial and fungal community composition differed significantly with compost application (Fig. 5C, D; PERMANOVA, bacteria: *p* = 0.001; fungi: *p* = 0.001). Bacterial community composition was also affected by microbiome origin and the compost × microbiome-origin interaction (Fig. 5C; *p* = 0.012 and *p* = 0.016, respectively), whereas these effects were not significant for fungal communities (Fig. 5D; *p* = 0.282 and *p* = 0.272, respectively). Community variability, as measured by beta dispersion, also differed among treatments for bacterial but not fungal communities (Fig. S2B). For bacteria, the effect of compost on variability depended on microbiome origin (Fig. S2B; compost x microbiome interaction, *p* = 7.06 × 10⁻⁴), with Farm 2 communities showing greater variability without compost than with compost. Compost application also reduced fungal community dispersion (Fig. S2B; compost main effect, *p* = 7.32 × 10⁻⁵), however there was no significant compost × microbiome-origin interaction (*p* = 0.94). These results indicate that compost altered both the composition and variability of rhizosphere microbial communities, whereas effects of microbiome origin were more evident for bacterial communities.

Differential abundance analysis identified 41 bacterial genera in the rhizosphere that responded significantly to compost application, including 14 that increased and 27 that decreased in relative abundance (Fig. 6A; Table S4; *p* < 0.05; log FC > 1.5). The bacterial genera showing the largest increases with compost application were *Sphaerobacter* (log FC = 5.53), *Litorilinea* (log FC = 5.07), and *Lysobacter* (log FC = 3.75). The largest decreases were observed for *Massilia* (log FC = −4.98), *Oxalophagus* (log FC = −4.95), and *Rhizobium* (log FC = −4.51) (Table S4). In contrast, only six fungal genera in the rhizosphere changed in relative abundance in response to compost application (Fig. 5A; Table S4; *p* < 0.05; log FC > 1.5). *Pseudogymnoascus* (log FC = 6.16), *Mortierella* (log FC = 4.24), and *Arachniotus* (log FC = 3.15) increased in relative abundance, whereas *Trichoderma* (log FC = −5.33), *Penicillium* (log FC = −4.29), and *Rhodotorula* (log FC = −3.98) decreased (Fig. 6A; Table S4). Microbiome origin was also associated with differences in specific bacterial taxa. Thirteen bacterial taxa differed significantly between plant rhizospheres of plants grown in the microbiome from Farm 1 and Farm 2 (Fig. 6B; Table S5). Taxa with the highest relative abundance in Farm 1 conditioned rhizospheres included *Paenibacillus kobensis* (log FC = 3.39), *Singulisphaera acidiphila* (log FC = 3.27), and *Niallia circulans* (log FC = 3.06), and in Farm 2 conditioned rhizospheres included *Asticcacaulis benevestitus* (log FC = 3.25), *Devosia soli* (log FC = 3.18), and *Rhodanobacter fulvus* (log FC = 2.10). No fungal taxa differed significantly in the rhizosphere of plants grown in the microbiome from Farm 1 and 2 (Table S5).

**Fig. 6.**
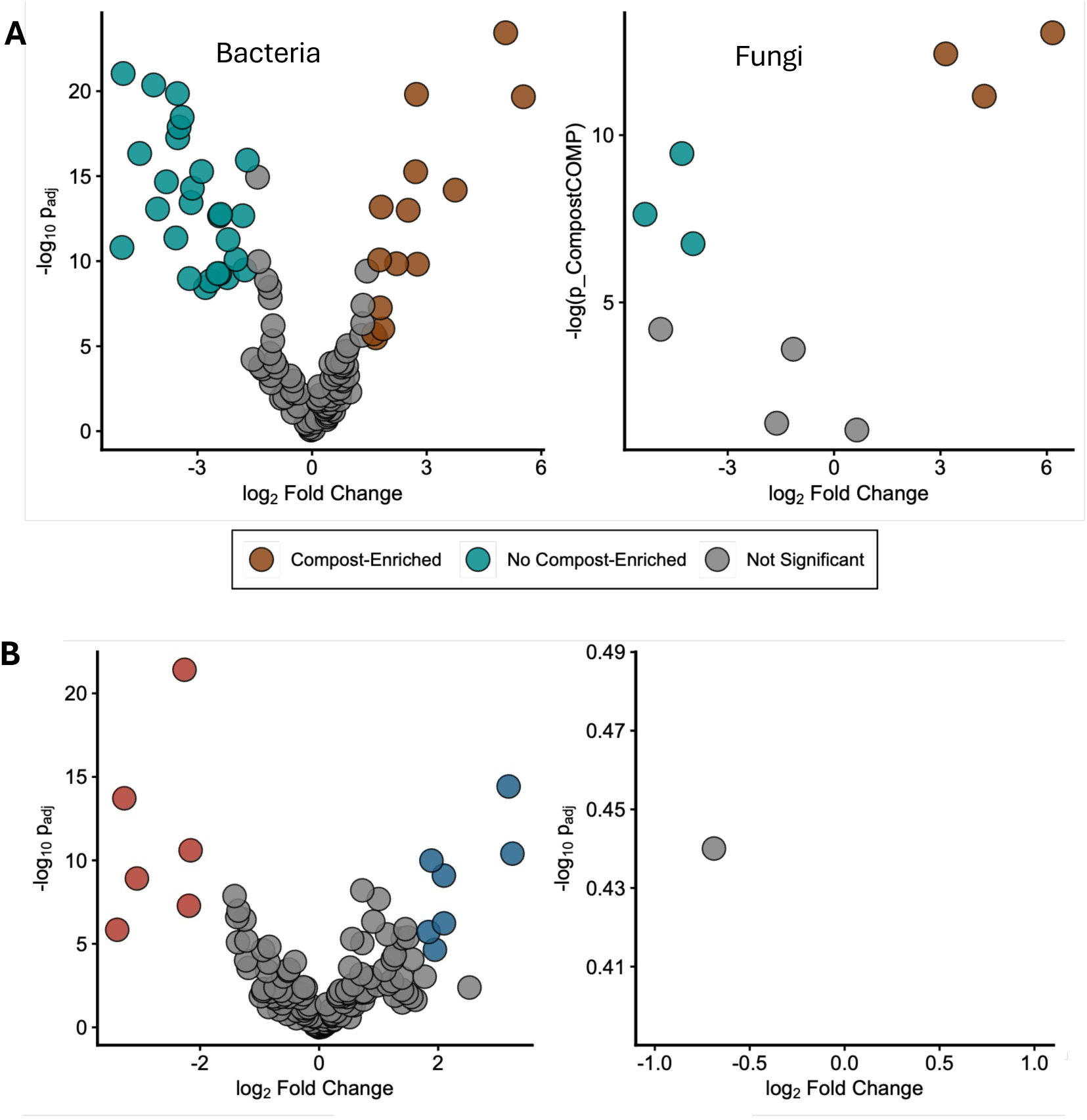
Compost application and microbiome origin impacts microbiome composition. A volcano plot was used to illustrate differentially abundant bacteria and fungi (B) enriched from compost application. N = 14. Microbes that significantly change from an ANCOMBC test with holm-adjusted-p < 0.01 are plotted by color: blue (enriched in no compost application) or brown (enriched in compost application). Those taxa that did not significantly change are colored gray. N = 8. A volcano plot was used to illustrate differentially abundant bacteria and fungi (B) enriched Farm 1 compared to Farm 2. N = 14. Microbes that significantly change from an ANCOMBC test with holm-adjusted-p < 0.01 are plotted by color: red (enriched in Farm 1 slurry) or blue (enriched in Farm 2 slurry). Those taxa that did not significantly change are colored gray. N = 8.

## Discussion

Organic agricultural systems often support greater soil microbial abundance compared to conventionally managed systems [53]. Understanding the mechanisms driving this difference is important because soil microbes can influence plant nutrient acquisition, growth, and resilience to biotic and abiotic stress [54–56]. Compost amendments, which are commonly applied in organic agriculture [57], may, in part, contribute to functional differences in the microbiomes of organic and conventional agricultural systems. In this study, compost application altered the relative abundance of specific microbial taxa across working organic farms (Fig. 2) and substantially restructured rhizosphere microbial communities under controlled laboratory conditions (Fig. 5, 6)) but did not alter microbiome-mediated plant-aphid dynamics (Fig. 4). Instead, soil microbiome origin was the strongest determinant of variation in aphid populations and defense-related metabolites (Fig. 4). Thus, measurable changes in microbial diversity and community composition do not always translate into corresponding changes in microbiome-mediated plant resistance. While increased biodiversity is often positively associated with ecosystem functions and productivity [58], our findings highlight that higher diversity is not a universal predictor for microbiome functions in plant-insect interactions or plant defense (Fig. 4, Fig. 5). More broadly, these findings suggest that the functional consequences of agricultural management may depend strongly on the identity and ecological context of resident microbial communities, rather than on microbial diversity alone.

Our findings are consistent with Badri et al. [59] who found that feeding by cabbage looper (*Trichoplusia ni*) larvae was reduced on Arabidopsis roots when grown in certain soil microbiomes extracts. Similarly, we found that kale plants grown with the microbiome from Farm 4 (yellow) supported lower aphid populations and SA concentrations than plants grown with microbiomes from Farms 1 (red) and 2 (blue) (Fig. 4). Rhizosphere microbial communities also differed between plants grown with the Farm 1 and Farm 2 microbiomes (Fig. 5,6). SA is often antagonistically regulated by jasmonic acid (JA), an important regulator of plant defenses against many herbivorous insects. Thus, plants grown with the Farm 4 microbiome may have had greater JA-associated defense responses, which could contribute to both reduced SA concentrations and decreased aphid populations, however, JA was not measured in this study. The soil samples from our study originated from farms across New York State, and regional biogeography has been identified as an important predictor of rhizosphere microbiome structure and function [60,61]. Furthermore, Lupatini, et al. [15] reported greater bacterial community variability across organically managed soils relative to conventional systems. Since our soil samples came from different organically managed farms, site-specific variation in resident microbial communities may have contributed to functional variation observed among microbiome origins.

Plant defense-related functions of the soil microbiome can depend on the presence of specific keystone taxa or functional guilds [62]. Some rhizosphere microbes have been shown to promote Induced Systemic Resistance (ISR) in plants [63] through the production of microbe-associated peptides and proteins (e.g., flagellin, cellulose-degrading proteins [64]) or small molecules (e.g., siderophore, volatile [65,66]). In a recent study, a structural equation modeling approach was used to identify microbiome changes in bulk soil that are associated with specific organic practices and functions in plant defense, and increases in SA were associated with a reduced abundance of *Paenibacillus senegalensis* [29]. Conversely, in our study, two *Paenibacillus* taxa (*P. kobensis, P. glycanilyticus*) increased in abundance in the rhizosphere of plants with the highest SA concentrations (Table S5; Fig. 4). While we did not detect any significant change in *P. senegalensis,* the contrasting associations among members from the same genus highlight the potential for species-specific relationships between microbial taxa and plants or differences in bulk and rhizosphere soils. Overall, these findings further suggest soil health assessments and interventions aimed at boosting plant defense may benefit from focusing on specific keystone taxa rather than general diversity.

The presence of nutrient-cycling bacteria in the rhizosphere can support plant growth and soil health by increasing nutrient availability and other soil biological functions [67], and has been associated with reduced abundance of insect herbivores [68]. Indeed, when compared to Farm 2 (blue), the rhizosphere from plants grown in Farm 1 (red) microbiomes were enriched in several potentially plant-beneficial taxa, including *Niallia circulans* (formerly *Bacillus circulans*) (Table S5)*. N. circulans* has potassium (K) mobilizing capabilities [69,70], and has been investigated as a K-solubilizing inoculant for agricultural crops [71]. Potassium-solubilizing microorganisms increase soil K bioavailability [72], a parameter that has been associated with both aphid populations [73] and SA signaling [74]. For example, aphid abundance was higher on soybean [73] and canola (*Brassica napus*) [75] grown in K deficiency than K sufficiency. One potential mechanism may be related to changes in phloem asparagine, an important source of nitrogen for aphids, which has been reported to decrease with increasing soil K concentration [73]. Although not significant, aphid populations were lower on plants grown with the Farm 1 microbiome treatment compared to Farm 2 (Fig. 4). Together, these observations raise the possibility that enrichment of K-mobilizing taxa such as *N. circulans* could contribute to microbiome-dependent differences in plant nutritional status and defense, however this mechanism remains to be tested directly.

While compost substantially altered the rhizosphere microbiome in the laboratory trials (Fig. 5,6), consistent with previous reports [76], no effect on aphid populations was observed (Fig. 4A). Compost has been shown to have variable effects on plant-insect dynamics, with some reports indicating suppressive [e.g., potato aphid (*Macrosiphum euphorbiae*) in tomato [77]] and others suggesting no effect [e.g., flea beetle (*Epitrix fuscula*) in eggplant [78])]. These differences across studies may be partially a result of the different compost formulations used. Mdellel et al. [79], for example, found that green peach aphid growth and populations on pepper were dependent on the type of compost applied, and suggested differences may be due to differences in compost N concentrations. Many of the bacteria that increased with compost application in our laboratory trial belonged to Actinomycetota or Bacillota, phyla commonly associated with rhizosphere interactions, organic matter decomposition, and nutrient cycling [80–86]. For example, *Gaiella,* a member of Actinomycetota, was enriched by compost in our study (Fig. 6A, Table S4). Numerous Ascomycota fungi have been shown to be dominant under composted conditions [87–90]. Within this group, the *Pseudogymnoascus* genus was enriched in the compost treatments in our study (Fig. 6A; Table S4) and has been associated with nutrient cycling in soil environments [91]. Compost application introduces microbial taxa while also providing organic substrates that alter resource availability and microbial interactions [21,92]. Indeed, Liu et al. [93] determined that microbiome connectivity increased after compost application by examining co-occurrence networks and identified connectivity as an indicator of this increased functionality. Although microbial connectivity was not measured in our study, the strong compositional response to compost suggests that changes in microbial interactions and functional organization could represent an important avenue for future investigation.

Our results demonstrate that increased soil microbial diversity and composition shifts that are often associated with organic farms and practices do not always translate into enhanced microbiome-mediated functions in plant defense. However, microbiomes originating from different farms altered both aphid populations (Fig. 4A) and plant chemistry (Fig. 4B), further supporting the role of the soil microbiome in bottom-up control of insect pests [94]. Taken together, these findings highlight the need for additional studies examining microbiome functional composition and the roles of specific microbial taxa or guilds in plant defense outcomes. Furthermore, it is well known plants select for specific microbes through root exudates [95,96], the mechanisms governing recruitment or persistence of defense-associated soil microbes remain poorly understood. In our study compost application had a marked effect on rhizosphere bacterial and fungal communities in laboratory conditions (Fig. 5), however, in field condition compost application did not alter the soil microbiome (Fig. 3). Biotic and abiotic legacies in the field strongly affect the degree to which compost modulates the soil microbiome [97] and differences in the microbiome outcomes between greenhouse and field trials and bulk and rhizosphere soils have been observed [98]. Future efforts to harness soil microbiomes for pest suppression may therefore benefit from focusing not simply on increasing microbial diversity, but on identifying the community contexts and functional taxa that consistently support plant resistance.

## List of abbreviations

PGPR: Plant growth promoting rhizobacteria
ISR: Induced systemic resistance
SA: Salicylic acid
PCoA: Principal coordinate analysis
LSD: Least Significant Difference
HSD: Honest Standard Difference
PERMANOVA: Permutational Multivariate Analysis of Variance
OTUs: Organizational taxonomic units
K: Potassium

## Declarations

### Ethics approval and consent to participate

Not applicable

### Consent for publication

Not applicable

### Availability of data and materials

Data and code corresponding to the analyses done in this manuscript can be found at https://github.com/marydixon/microbiome_compost_aphid.

Sequencing data can be found in the NCBI sequence read archive (BioProject No. PRJNA1468248).

### Author’s Contribution

EJM, CLC conceptualization; EJM, CLC, EHB methodology; MMD, EJM formal analysis; EJM, BC investigation; MMD, EJM, BC, CLC writing with feedback from all authors; MMD, EJM visualization; CLC supervision. All authors read and approved the final manuscript

## Acknowledgements

We thank Dr. Dinesh Pujara for his assistance in running samples on the LC/MS for this study. This work was supported in part by a USDA ORG Grant to C. L. C. (#2022-51106-38007), a NIFA-USDA Hatch Multistate Grant to C. L. C. (#7000327), a NIFA-USDA Hatch Grant to C. L. C. (#7003512), and start-up funds to C. L. C. The findings and conclusions in this publication have not been formally disseminated by the U.S. Department of Agriculture and should not be construed to represent any agency determination or policy.

---

**Supplemental Figure 1.**
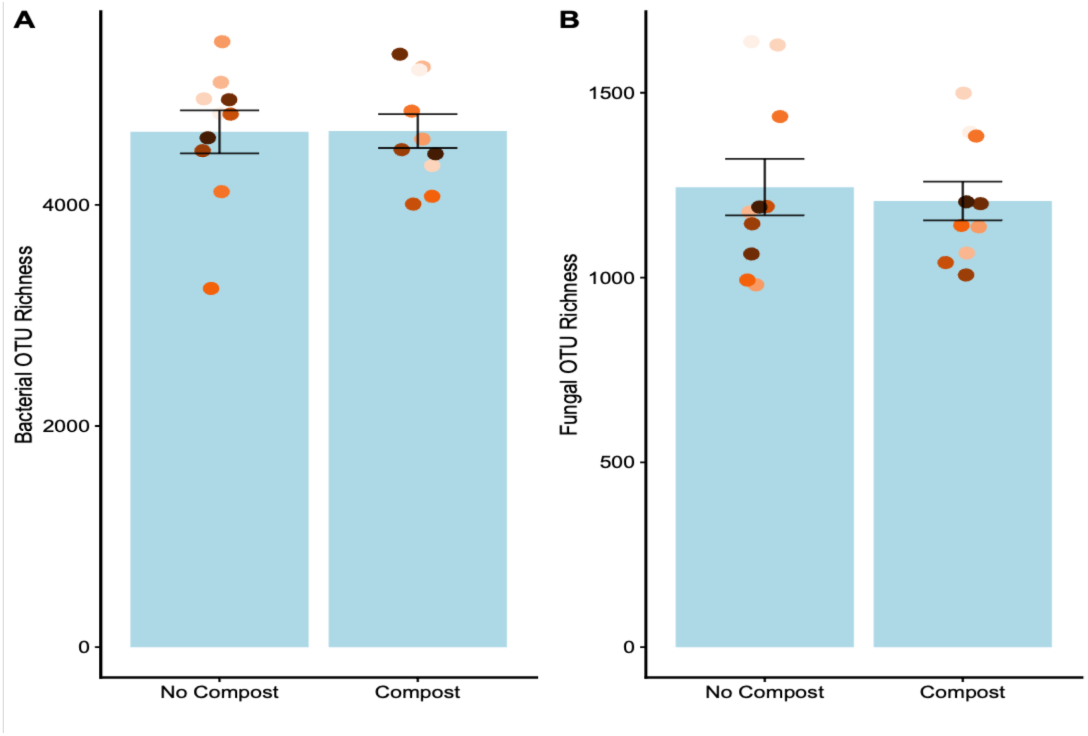
OTU richness and Simpson’s diversity for bacterial and fungal taxa in field samples collected from Bloom et al. (2024).

**Supplemental Fig. 2.**
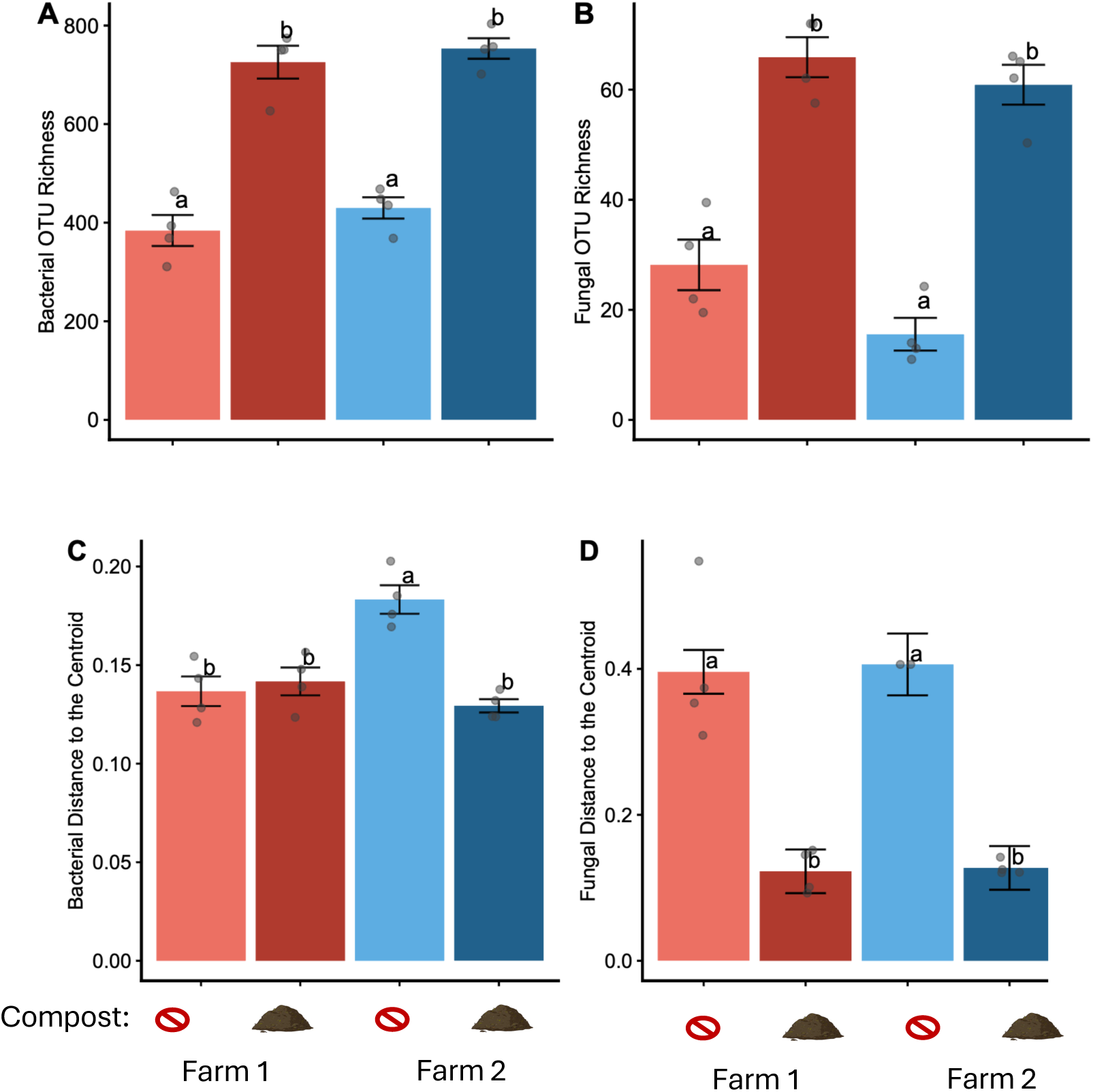
Richness was measured for bacteria (A) and fungi (B) in no compost and composted plots.Beta dispersion was calculated for (C) bacteria and (D) fungi . An ANOVA was run with Tukey HSD for pairwise comparisons. Different letters denote differences at α = 0.05 for the farm site used for soil collection. Bars represent mean ± SE. Color denotes the farm origin of the soil used and the compost application: light red and dark red (Farm 1) & light blue and dark blue (Farm 2). N = 16.

